## supplemental methods for "Habitual exercise evokes fast and persistent adaptation during split-belt walking"

### Supplemental Statistical Methods

For each of the four work rate outcome measures, we built-up one-exponent models — assuming that adaptation of the outcome measures occurs over a single timescale — that did not include a fixed effect for group (eq. S1) and that did include a fixed effect for group (eq. S2). Here, *c* is the estimated plateau in work rate if strides went to infinity; *a* is the initial value of work rate; and *r* is the growth rate of work rate adaptation. In the models that included a fixed effect for group, all three model parameters (*c*, *a*, *r*) were allowed to differ for MOVE vs. notMOVE.

$$Work\ rate = c + a * e^{-\frac{stride}{r}} + (c|ID) \qquad \text{S1}$$

$$Work\ rate = \left(a * e^{-\frac{stride}{r}} + c\right) \sim Group + (c|ID) \qquad \text{S2}$$

Then, for each of the work rate outcome measures and for SLA, we built-up two-exponent models — assuming that adaptation of the outcome measures occurs over two timescales — that did not include a fixed effect for group (eqs. S3 and S5) and that did include a fixed effect for group (eqs. S4 and S6). Here, *c* is the estimated plateau if strides went to infinity; $a_f$ is the initial value of the fast component of adaptation; $r_f$ is the growth rate of the fast component of adaptation; $a_s$ is the initial value of the slow component of adaptation; $r_s$ is the growth rate of the slow component of adaptation. In the models that included a fixed effect for group, all five model parameters ($c$, $a_f$, $r_f$, $a_s$, $r_s$) were allowed to differ for MOVE vs. notMOVE.

$$SLA = c + a_f * e^{-\frac{step}{r_f}} + a_s * e^{-\frac{step}{r_s}} + (c|ID) \qquad \text{S3}$$

$$SLA = (c + a_f * e^{-\frac{step}{r_f}} + a_s * e^{-\frac{step}{r_s}}) \sim Group + (c|ID) \qquad \text{S4}$$

$$Work\ rate = c + a_f * e^{-\frac{stride}{r_f}} + a_s * e^{-\frac{stride}{r_s}} + (c|ID) \qquad \text{S5}$$

$$Work = \left( c + a_f * e^{-\frac{stride}{r_f}} + \ a_s * e^{-\frac{stride}{r_s}} \right) \sim Group + (c|ID) \qquad \text{S6}$$

All models were fit with a maximum likelihood estimation and all models contained a random effect such that the outcome measure's estimated plateau was allowed to vary by participant.
