## supplemental table 1 for "Habitual exercise evokes fast and persistent adaptation during split-belt walking"

**Table S1.** Fits of mixed effects linear models for each outcome measure.

| *Variable* | *1-exponent model* | *1-exponent + group model* | *2- exponent model* | ***2-exponent + group model*** |
|---|---|---|---|---|
| **Step length asymmetry** | | | | |
| AIC | — | — | -105693.6 | -106168.7 |
| BIC | — | — | -105636.3 | -106070.6 |
| Log-likelihood | — | — | 52853.78 | 53096.36 |
| Number of observations | — | — | 26336 | 26336 |
| Number of participants | — | — | 32 | 32 |
| SD: ID (*c*) | — | — | 0.038 | 0.038 |
| SD: Residual (*c*) | — | — | 0.032 | 0.032 |
| **Positive work rate of the fast leg** | | | | |
| AIC | 2409.80 | 2230.21 | 2211.81 | 2021.38 |
| BIC | 2447.25 | 2290.13 | 2264.23 | 2111.25 |
| Log-likelihood | -1199.90 | -1107.11 | -1098.90 | -998.69 |
| Number of observations | 13216 | 13216 | 13216 | 13216 |
| Number of participants | 32 | 32 | 32 | 32 |
| SD: ID (*c*) | 0.441 | 0.435 | 0.441 | 0.435 |
| SD: Residual (*c*) | 0.263 | 0.261 | 0.261 | 0.259 |
| **Negative work rate of the fast leg** | | | | |
| AIC | -12275.95 | -12209.38 | -12285.29 | -12350.91 |
| BIC | -12238.51 | -12149.46 | -12232.87 | -12261.04 |
| Log-likelihood | 6142.98 | 6112.69 | 6149.65 | 6187.46 |
| Number of observations | 13216 | 13216 | 13216 | 13216 |
| Number of participants | 32 | 32 | 32 | 32 |
| SD: ID (*c*) | 0.352 | 0.347 | 0.352 | 0.347 |
| SD: Residual (*c*) | 0.151 | 0.151 | 0.151 | 0.150 |
| **Positive work rate of the slow leg** | | | | |
| AIC | -12371.19 | -12699.95 | -12624.73 | -12838.49 |
| BIC | -12333.74 | -12640.04 | -12572.31 | -12748.6 |
| Log-likelihood | 6190.59 | 6357.98 | 6319.37 | 6431.24 |
| Number of observations | 13216 | 13216 | 13216 | 13216 |
| Number of participants | 32 | 32 | 32 | 32 |
| SD: ID (*c*) | 0.232 | 0.231 | 0.232 | 0.231 |
| SD: Residual (*c*) | 0.150 | 0.148 | 0.149 | 0.147 |
| **Negative work rate of the slow leg** | | | | |
| AIC | 5319.77 | 5203.29 | 4962.98 | 4848.64 |
| BIC | 5357.22 | 5263.21 | 5015.40 | 4938.51 |
| Log-likelihood | -2654.89 | -2593.65 | -2474.49 | -2412.32 |
| Number of observations | 13216 | 13216 | 13216 | 13216 |
| Number of participants | 32 | 32 | 32 | 32 |
| SD: ID (*c*) | 0.579 | 0.572 | 0.579 | 0.572 |
| SD: Residual (*c*) | 0.293 | 0.292 | 0.289 | 0.288 |
