## Supplementary figures and images for "Habitual exercise evokes fast and persistent adaptation during split-belt walking"

### supplemental figure 1

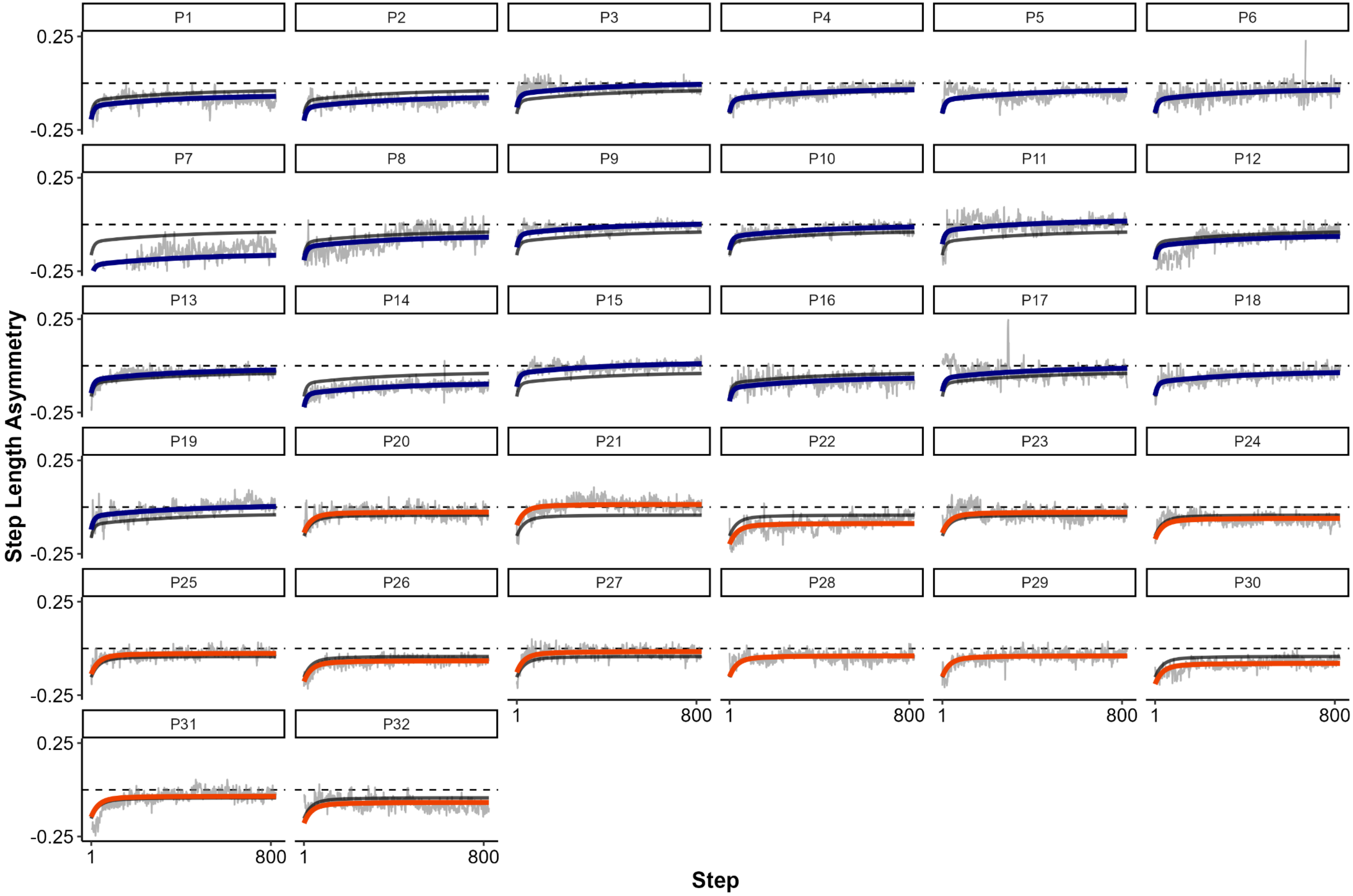

### supplemental figure 2

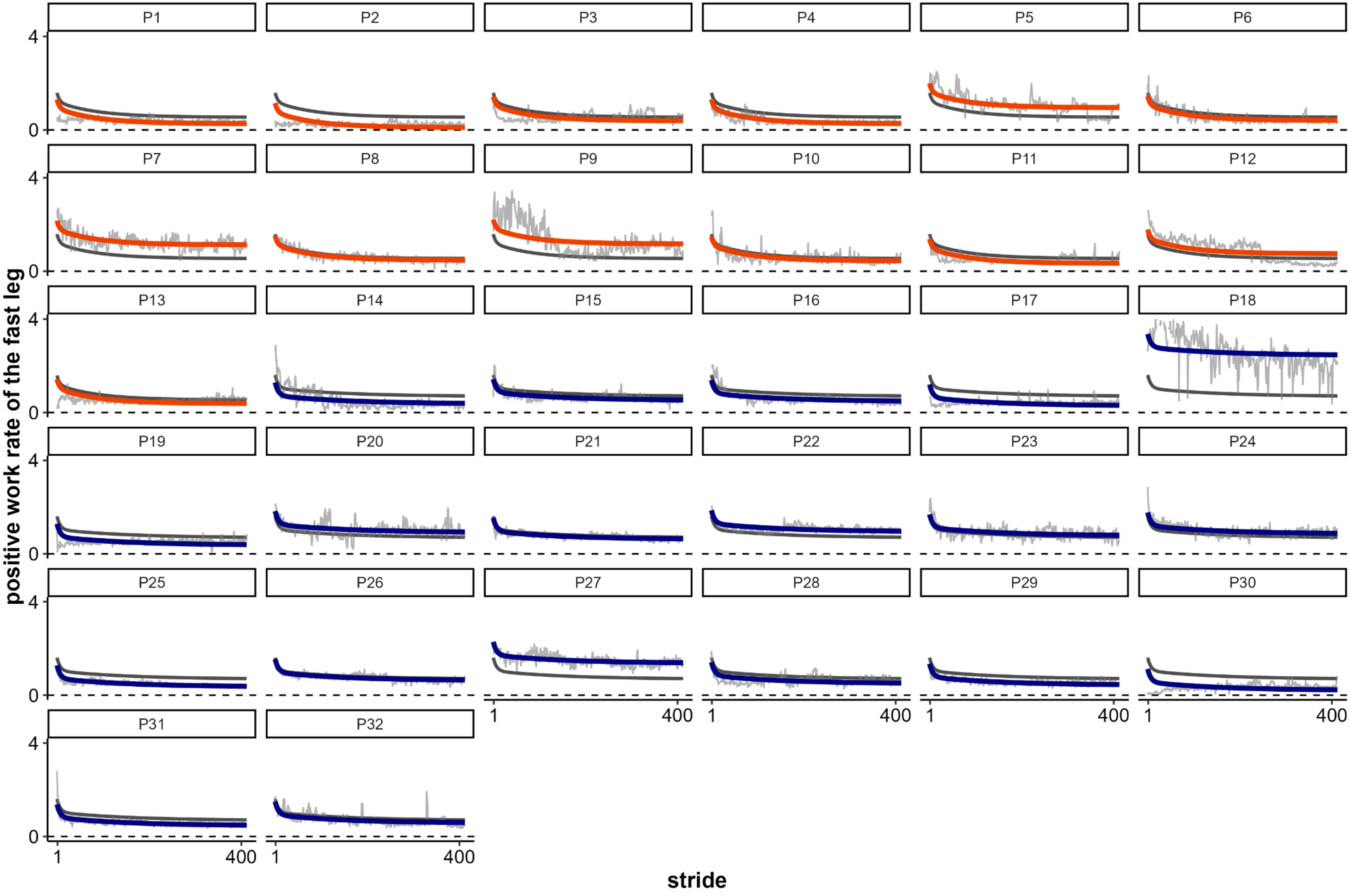

### supplemental figure 3

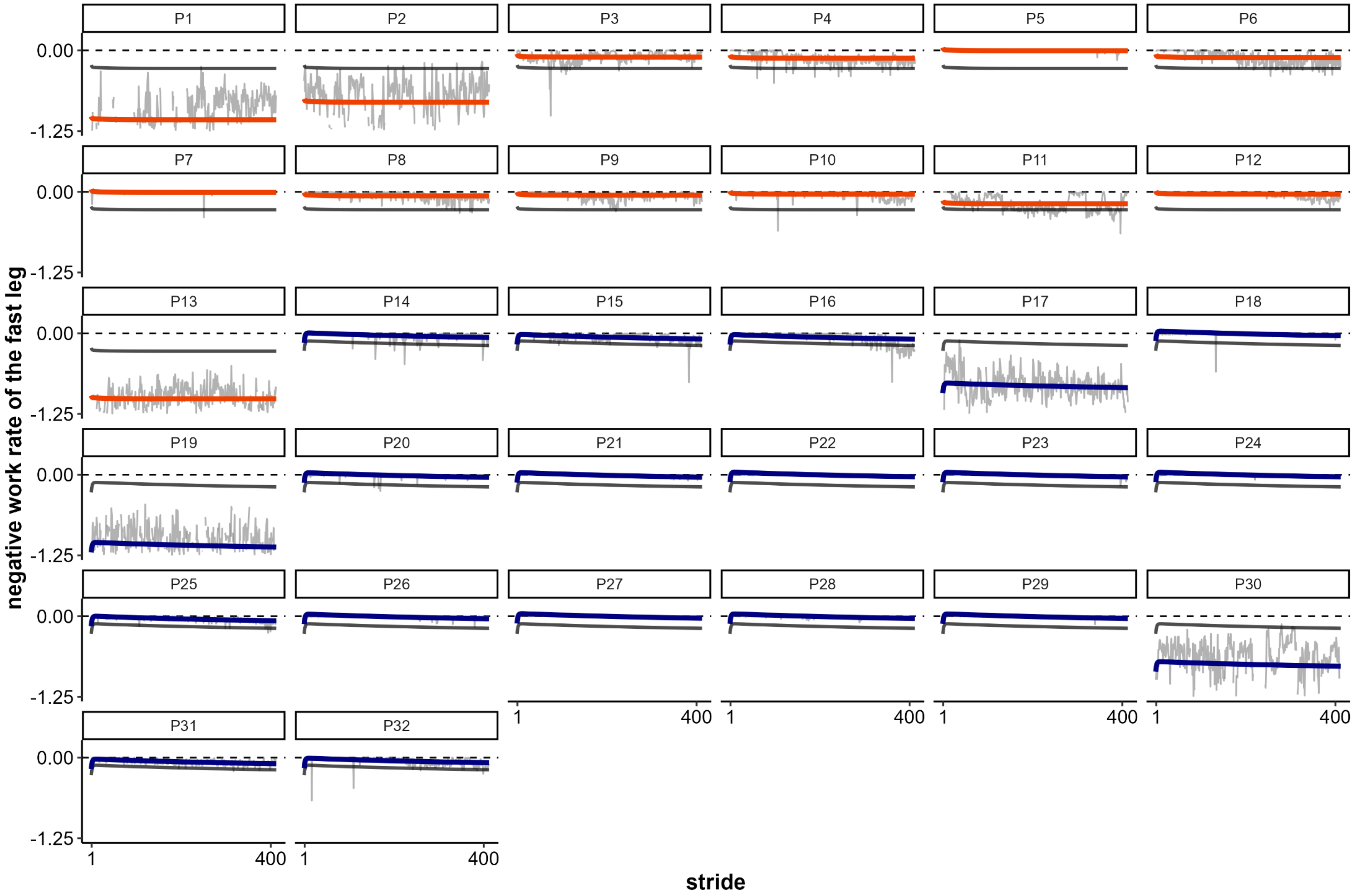

### supplemental figure 4

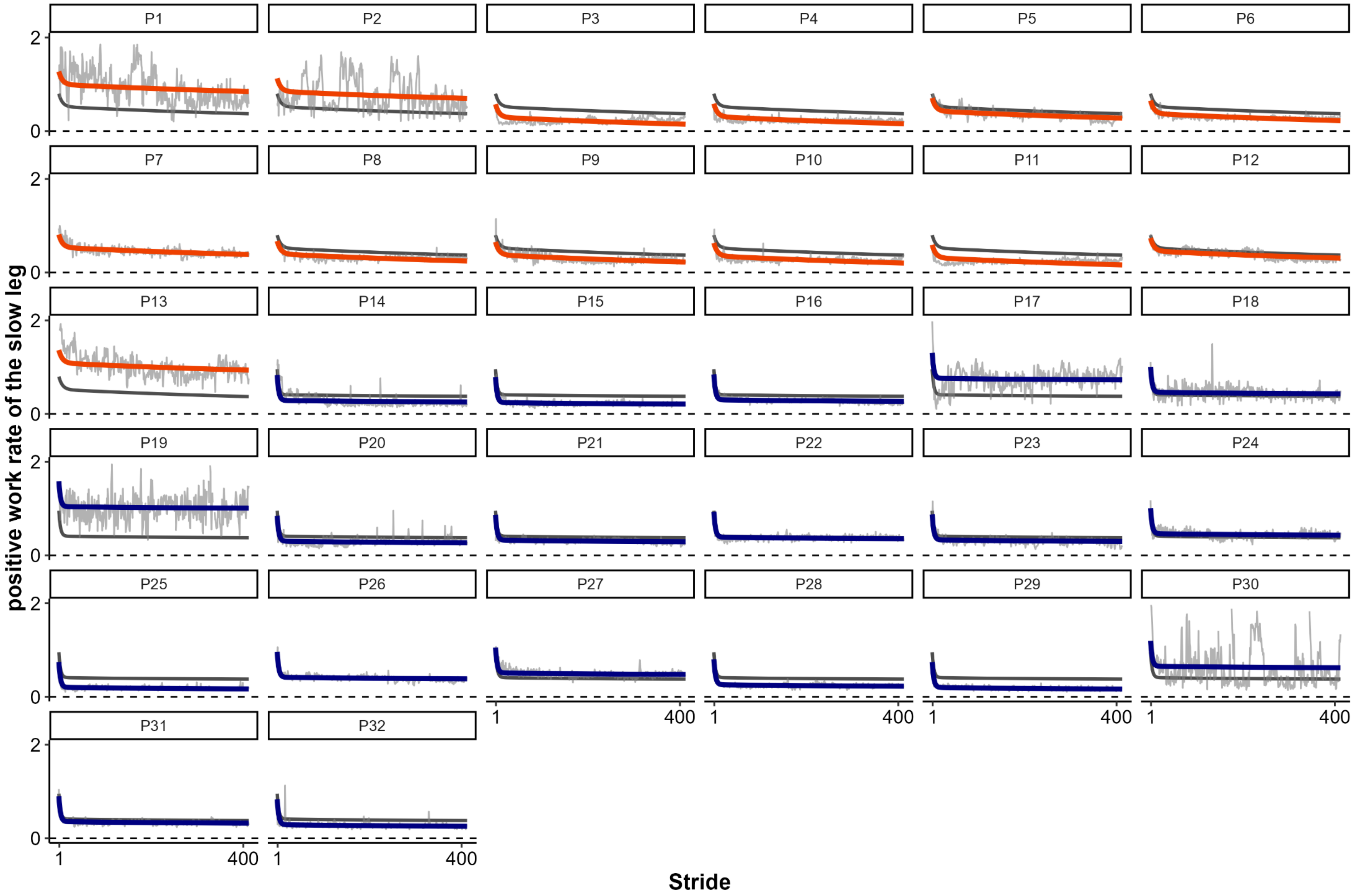

### supplemental figure 5

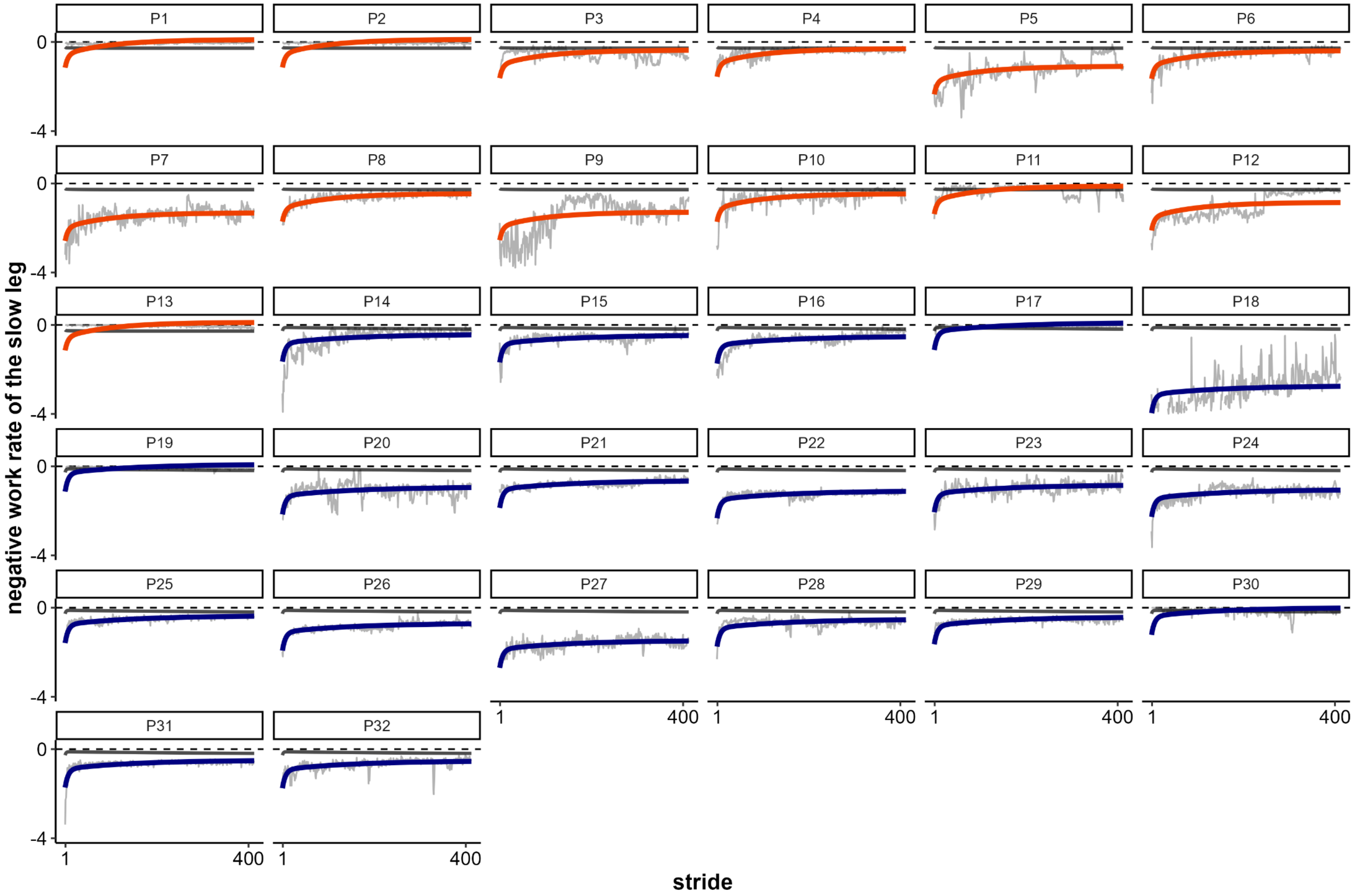
